# DNA-based delivery of incretin receptor agonists using MYO Technology leads to durable weight loss in a diet-induced obesity model

**DOI:** 10.1101/2025.05.30.656889

**Authors:** Linda Sasset, Andrew Cameron, Carleigh Sussman, Luisa Rubinelli, Debnath Maji, Robert Miller, Andy Thompson, Delcora Campbell, Melanie R. Walker, Marek M. Drozdz, Rachel A Liberatore

## Abstract

Therapeutic proteins have seen a substantial increase in clinical development and use across many disease areas. Despite their broad applicability, significant drawbacks limit access to many of these drugs, including: i) high manufacturing costs; ii) administration via time-consuming infusions; iii) frequent dosing, sometimes even daily; and iv) requirements for low temperature storage.

MYO Technology was developed to overcome these barriers. The MYO Technology platform consists of therapeutic-encoding plasmid DNA (pDNA), and a proprietary medical device for intramuscular injection and delivery of electrical pulses. These pulses enable the *in vivo* electroporation of muscle cells and uptake of injected pDNA, leading to the production, secretion, and delivery of the therapeutic protein into peripheral circulation. MYO Technology offers several advantages over standard delivery of therapeutic proteins; pDNA manufacturing is a simpler and less specialized process compared to protein manufacturing, and pDNA is very stable and lacks most cold chain requirements. Furthermore, administration using MYO Technology takes only a few minutes, and the serum level of a therapeutic protein can potentially be maintained for many months without the need for redosing.

Incretin receptor agonists (IRAs) are a class of therapeutic proteins that have recently come to prominence as powerful weight and glucose control drugs, and are used for the treatment of type 2 diabetes (T2D) and obesity. Semaglutide and tirzepatide, currently the most widely used within this class, are both potent molecules, but have a short half-life, requiring weekly administration by subcutaneous injections. Moreover, since their clinical benefits rapidly disappear upon treatment cessation, T2D and obese patients may have a life-long dependency on IRAs, and the requirement for weekly injections can negatively affect the quality of life and the adherence to therapy, as well as create a significant financial burden. Therefore, increasing the interval between injections has become one of the major goals in the field.

Here, we present our preclinical studies on the delivery of IRAs with MYO Technology. Animal proof-of-concept studies demonstrate that MYO Technology-delivered IRAs are functional, and efficacious in promoting long-lasting weight and glucose control in mouse models of diet-induced obesity. Moreover, engineering the IRAs to facilitate blood–brain barrier penetration further enhances treatment efficacy, with benefits persisting beyond one year following a single administration. Together, these findings highlight MYO Technology’s potential to transform care for patients with T2D and obesity by enabling long-lasting therapeutic effects with minimal dosing, ultimately improving quality of life and treatment adherence.

## INTRODUCTION

Glucagon like peptide-1 (GLP-1) is an emerging glucose control and weight loss drug, which is used in the treatment of type 2 diabetes (T2D) and obesity. This peptide is administered subcutaneously to patients, and upon binding to its receptor in different organs, GLP-1 exerts several effects: in the pancreas it enhances insulin secretion and inhibits glucagon release, in the stomach it delays gastric emptying, and in the hypothalamus it reduces hunger, increases the feeling of satiety, and alters feeding behavior, overall decreasing body weight and ameliorating metabolic dysregulation^1^.

A significant downside of GLP-1 is its short half-life, which is about 2-5 minutes in vivo^2^. Despite significant progress made in the field to incorporate modifications capable of improving its durability, the current GLP-1 class of drugs used in the clinic still has a half-life of only about one week, making weekly dosing necessary to maintain its therapeutic effects^3^.

Several molecules are currently approved for both obesity and T2D: liraglutide (Saxenda) and semaglutide (Ozempic/Wegovy) are GLP-1 receptor agonists, and tirzepatide (Mounjaro/Zepbound) is a double GLP-1 and GIP receptor agonist. All these molecules have an added fatty acid moiety, which binds albumin as a mechanism for half-life extension^4^. Some earlier generation molecules are only approved for T2D, such as dulaglutide (Trulicity), which incorporates an Fc fusion as a strategy for half-life extension^5^.

RenBio has evaluated intramuscular electroporation (IM-EP)-mediated delivery of plasmid DNA (pDNA), encoding GLP-1, using its MYO (Make Your Own) Technology, as a new therapeutic strategy. With this approach, GLP-1 is constantly produced for an extended period, overcoming the necessity for weekly dosing. This technology has been successfully employed in animal models of neutropenia for the delivery of granulocyte colony stimulating factor (G-CSF), another small therapeutic protein with a short (∼3.5 hours) half-life^6^.

MYO Technology takes advantage of the efficiency of in vivo transfection of skeletal muscle cells enabled by EP, and the durable protein production by those cells following transfection^7^. Indeed, expression of a monoclonal antibody or other therapeutic protein can be sustained for several months to a year or more, both in animal models, and most importantly, in a clinical setting^6^. Because of this durability, the potential exists to make a significant impact on the treatment of chronic diseases, including obesity^7^.

The GLP-1 class of drugs has recently transformed the treatment of obesity, though questions remain regarding the full suite of mechanisms of action of these molecules. In addition to direct action on the gastrointestinal tract, data suggest that targets in the brain are important for their activity^8^. Angiopep-2 is a peptide that has been shown to increase penetration of the blood-brain barrier (BBB) when fused to heterologous proteins^9^. Using a mouse model of diet-induced obesity (DIO), we demonstrate sustained weight loss and improved glucose metabolism for more than one year following a single dose of a GLP-1-Fc fusion protein with an appended Angiopep-2 moiety. This approach raises the prospect of dramatically reduced dosing frequency of this important class of therapeutics.

## RESULTS

### DNA-based delivery of GLP-1 peptide is efficacious in a diet induced obesity model

To evaluate the feasibility of DNA-based delivery of GLP-1 with MYO Technology we used a DIO mouse model^10–12^. Recombinant GLP-1 currently used in clinic has an added fatty acid moiety, which binds albumin and aids in half-life extension^4^ but this is incompatible with a DNA-based approach. We were able to alter the cleavage site recognized by DPP-4 in GLP-1, another strategy incorporated into the current class of GLP-1 drugs. DPP-4 is a protease that breaks down incretin hormones like GLP-1 and GIP, and plays a key role in the regulation of GLP-1 activity^13^.

To evaluate the effect of MYO-delivered GLP-1, mice were fed a high fat diet (HFD) for 8 weeks and then divided into four groups: two groups were kept on HFD and two groups switched to a standard diet (SD). We reasoned that the cohorts fed SD would better resemble a clinical scenario where obesity patients make dietary changes in addition to the drug treatment. Within each diet group, one subgroup was electroporated with the pDNA encoding GLP-1 and one kept as a control (Figure 1A). Mice that remained on HFD and received GLP-1 showed reduced weight gain in the first two weeks compared to the placebo controls (Figure 1B). This effect was transient, and after 2 weeks, all animals on HFD gained weight at the same rate. The therapeutic effect of MYO-delivered GLP-1 was much more pronounced in mice switched to SD (Figure 1C). The experimental cohort started losing weight immediately after IM-EP, showing over 15% weight loss compared to placebo control by week 2. Between week 3 and 5, however, animals started regaining weight, and by the end of the experiment, the placebo adjusted weight loss was at ∼5%.

**Figure 1.**
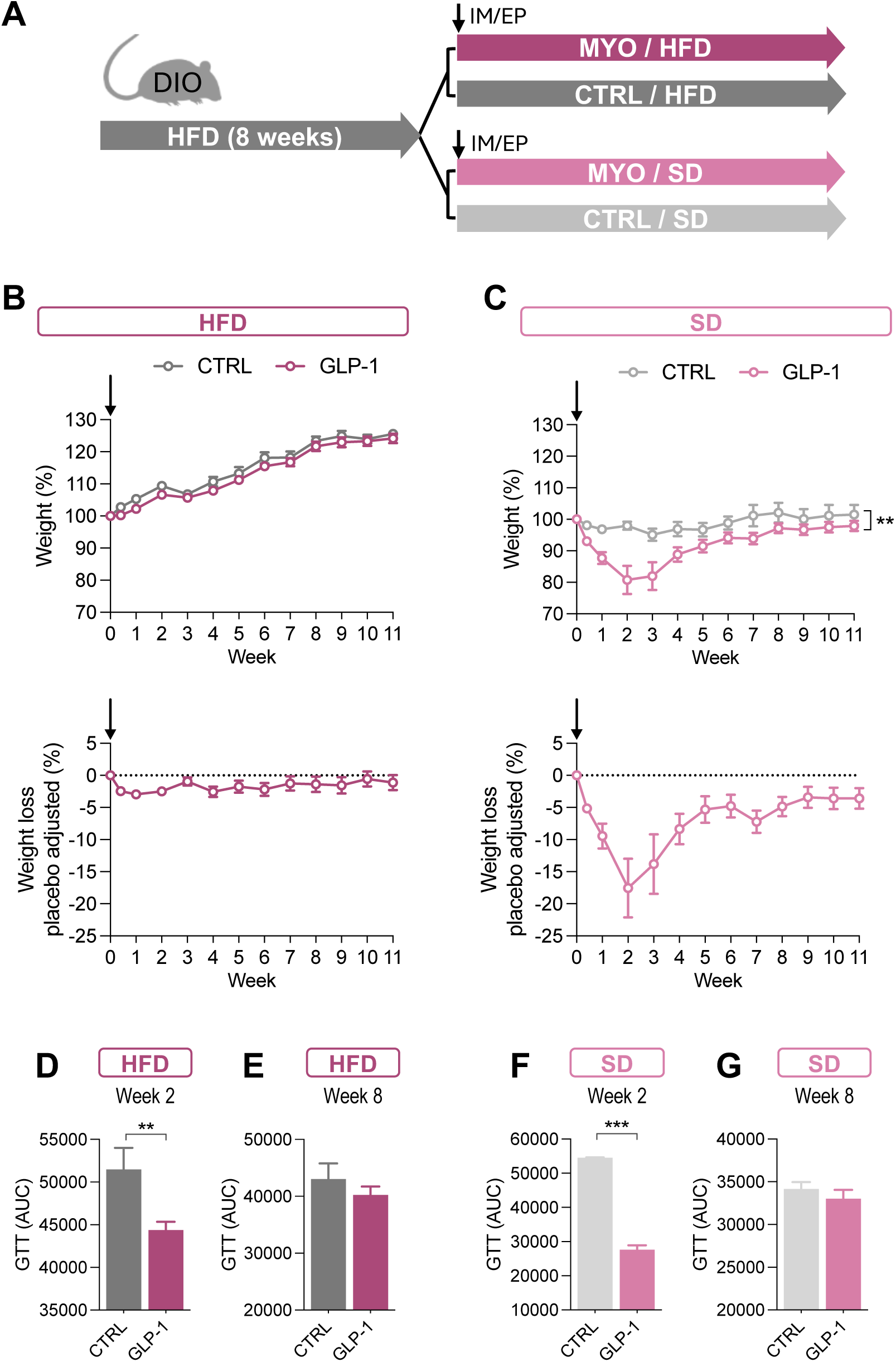
DNA-based delivery of GLP-1 peptide is efficacious in a diet induced obesity model. (A) Experimental design. Percentage of absolute (top panel) and placebo adjusted weight gain/loss (bottom panel) relative to the time of the EP (Week 0) in mice fed (B) high fat diet (HFD) (n=5), and (C) standard diet (SD) (n=5). Area under the curve (AUC) derived from glucose tolerance test (GTT) performed at (D) week 2, and (E) week 8 in mice fed HFD (n=5). AUC derived from GTT performed at (F) week 2, and (G) week 8 in mice fed SD (n=5). Data are expressed as mean ± SEM. ** p<0.01, *** p<0.01. Statistical significance was determined by Two-way mixed-effects ANOVA (B-C) and unpaired t-test (D-G).

Another important effect of GLP-1 molecules is the improvement of glycemic control. We therefore performed a glucose tolerance test (GTT) assay at different time points post IM-EP. GTT is a test that measures how the body regulates blood sugar levels after consuming a known amount of glucose^14^. Two weeks after IM-EP delivery, mice that received GLP-1 showed improved glucose clearance compared to controls in HFD groups (Figure 1D), demonstrating that the MYO-delivered GLP-1 is biologically active. We also examined glycemia at a later timepoint, when any weight loss benefit was already diminished. Not surprisingly, at eight weeks after IM-EP, we did not detect any difference in the glucose clearance rate (Figure 1E). The same was true for the animals switched to SD; two weeks after EP, animals that received GLP-1 showed improved GTT, but the effect had disappeared by week 8 (Figure 1F and G, respectively).

### GLP-1 fused to an Fc domain exhibits therapeutic effects comparable to those of semaglutide

Since the therapeutic effect of native GLP-1 peptide was limited in time, we decided to fuse GLP-1 with the sequence encoding an IgG Fc region in order to extend the molecule’s half-life and boost its bioavailability^5^. We focused on a murine Fc sequence to allow proof-of-concept studies in a DIO mouse model without evoking an anti-drug antibody response. The encoded protein is referred to as GLP-1-mIgG1(Fc) (Figure 2A).

**Figure 2.**
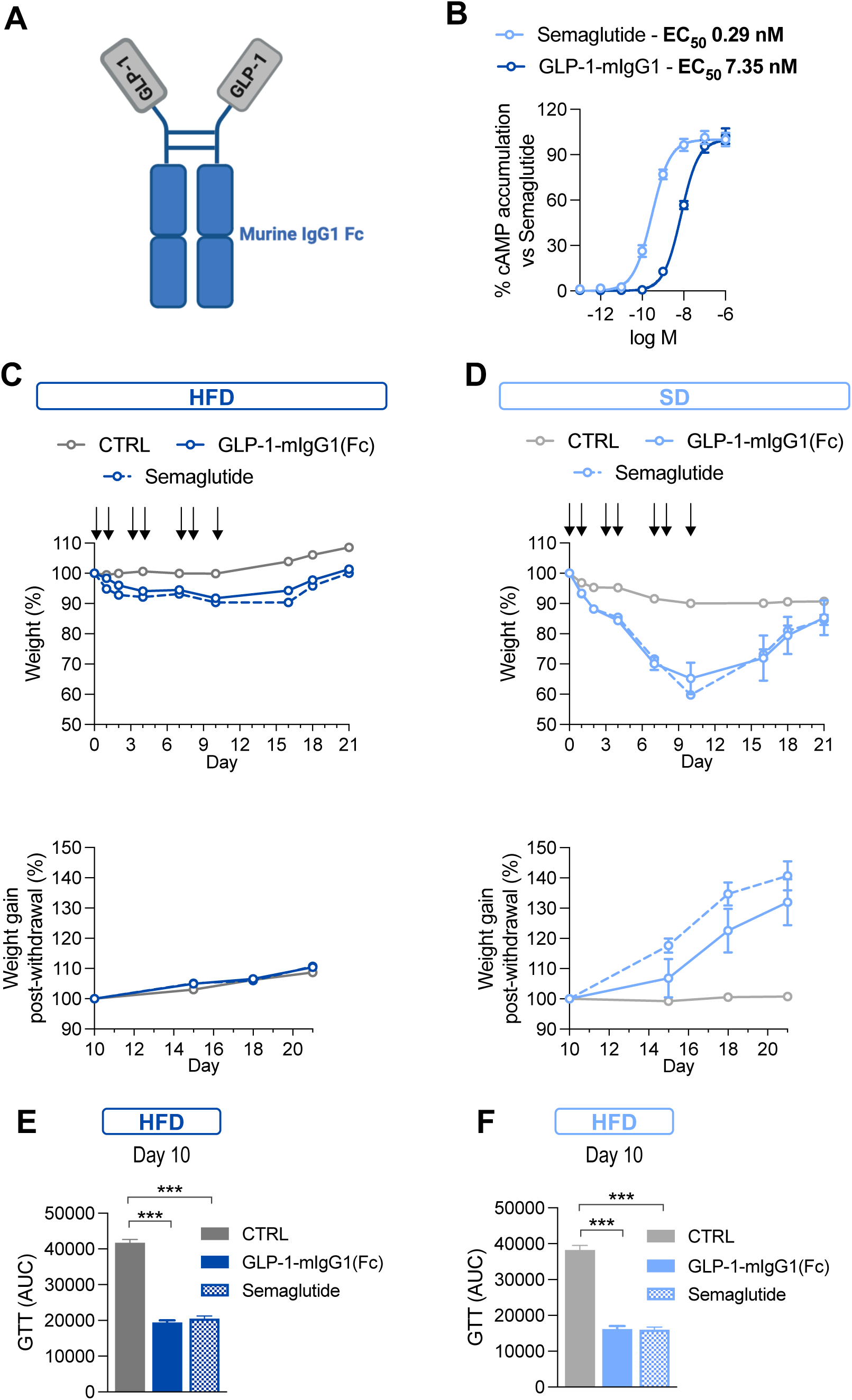
GLP-1 fused to an Fc domain exhibits therapeutic effects comparable to those of semaglutide. (A) Schematic of the protein. (B) Intracellular cAMP accumulation measured in HEK293T cells expressing GLP-1R. Data are expressed as % of cAMP relative to the highest semaglutide concentration (n=10 from 3 independent experiments). Percentage of absolute weight gain/loss relative to the time of the first injection (Day 0, top panel), and percentage of absolute weight gain relative to the time of the last injection (Day 10, bottom panel) in mice fed (C) HFD (n=5), and (D) SD (n=5). Area under the curve (AUC) derived from glucose tolerance test (GTT) performed at day 10 in mice fed (E) HFD (n=5), or (F) SD (n=5). Data are expressed as mean ± SEM. *** p<0.01. Statistical significance was determined by One-way ANOVA (E-F).

First, we compared the potency of GLP-1-mIgG1(Fc) with that of semaglutide. To do so, we developed an in vitro reporter assay^15^. We co-expressed GLP-1 receptor (GLP-1R) and a luciferase reporter under the control of a cAMP-responsive element (CRE) in HEK293T cells. GLP-1R activation leads to the activation of the adenylyl cyclase and consequent increase in cAMP generation which, in turn, activates CRE-dependent transcription. Using this assay, we evaluated GLP-1R activation by recombinant GLP-1-mIgG1(Fc) or semaglutide (Figure 2B). We determined semaglutide EC50 at 0.29 nM, which is comparable to 0.15 nM reported by Novo Nordisk in their FDA application (application #210913Orig1s000). GLP-1-mIgG1(Fc), albeit less potent than semaglutide, activated GLP-1R at low nanomolar ranges (EC50 = 7.35 nM).

Next, we evaluated GLP-1-mIgG1(Fc) compared to semaglutide in DIO mice when delivered as purified protein, instead of IM-EP of pDNA. We followed the same diet protocol described in Figure 1A. After randomization into HFD and SD groups, animals received either molecule subcutaneously at 30nmol/kg over a period of 10 days, then observed for another 11 days to monitor the effects of drug withdrawal.

In both HFD and SD groups, GLP-1-mIgG1(Fc) and semaglutide demonstrated comparable: i) reduction of body weight (Figure 2C and D, top panels); ii) weight rebound post-withdrawal (Figure 2C and D, bottom panels); and iii) improved glucose clearance (Figure 2E and F). These data confirm that the GLP-1 Fc fusion and semaglutide have comparable effects on weight and glucose metabolism in mice.

### DNA-based delivery of GLP-1 Fc fusion improves weight loss compared to native GLP-1

After confirming comparable potency of recombinant GLP-1-mIgG(Fc) and semaglutide, we delivered DNA-encoded GLP-1-mIgG1(Fc) with MYO Technology, following the same diet protocol as before (Figure 1A). Similar to the first study (Figure 1B), the cohort kept on a HFD lost minimal weight compared to controls (Figure 3A). The cohort switched to SD, however, started losing weight immediately after EP and showed much bigger weight loss than GLP-1 alone (Figure 3B). Importantly, even after animals started regaining weight past week 3, the placebo adjusted weight loss remained at ∼10% by the end of the experiment at week 6, which was a two-fold improvement from the GLP-1 alone. Assessment of the GTT showed a significant improvement in the glycemic control in groups that received GLP-1 Fc fusion. The difference was significant at 2 weeks after EP regardless of the diet and was maintained at week 6 in the SD cohort (Figure 3C-F).

**Figure 3.**
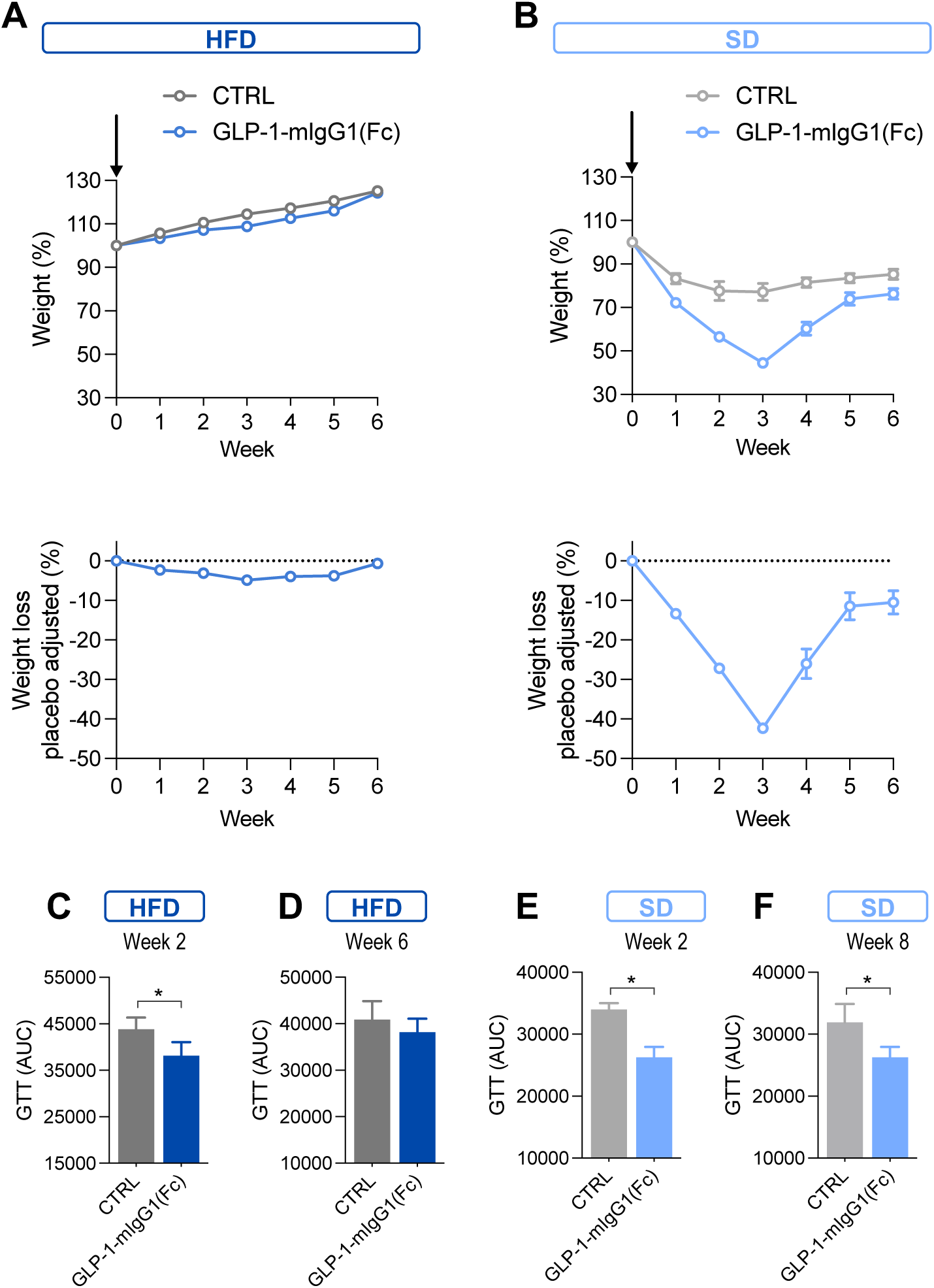
DNA-based delivery of GLP-1 Fc fusion improves weight loss compared to native GLP-1. Percentage of absolute (top panel) and placebo adjusted weight gain/loss (bottom panel) relative to the time of the EP (Week 0) in mice fed in mice fed (B) HFD (n=5), and (C) SD (n=5). Area under the curve (AUC) derived from glucose tolerance test (GTT) performed at (C) week 2, and (D) week 6 in mice fed HFD (n=5), and at (E) week 2, and (F) week 6 in mice fed SD (n=5). Data are expressed as mean ± SEM. * p<0.05. Statistical significance was determined by unpaired t-test (C-F).

### Enhanced BBB crossing dramatically improves GLP-1-mIgG1(Fc) efficacy

One of the mechanisms by which GLP-1 promotes weight loss is by binding to GLP-1R in the brain, particularly in the hypothalamus, and enhancing the feeling of satiety^1,16^. GLP-1-mIgG1(Fc) fusion is a relatively large protein and presumably unable to easily penetrate the BBB. The inability to pass through the BBB is an issue for several large molecules (e.g. antibodies) as well as for adeno-associated viral vectors, and several strategies have been recently developed to overcome this. One such strategy is the addition of Angiopep-2, a 19 amino acid peptide that binds to the low-density lipoprotein receptor related protein-1 (LRP-1) expressed by the endothelial cells of the BBB; Angiopep-2 binding to LRP-1 facilitates the transcytosis of the molecules conjugated to the peptide^17,18^. Therefore, we decided to assess whether the addition of Angiopep-2 to GLP-1-mIgG1(Fc) could improve the molecule’s efficacy in treatment of obesity. This new molecule is referred to as GLP-1-mIgG1(Fc)-Angp2 (Figure 4A).

**Figure 4.**
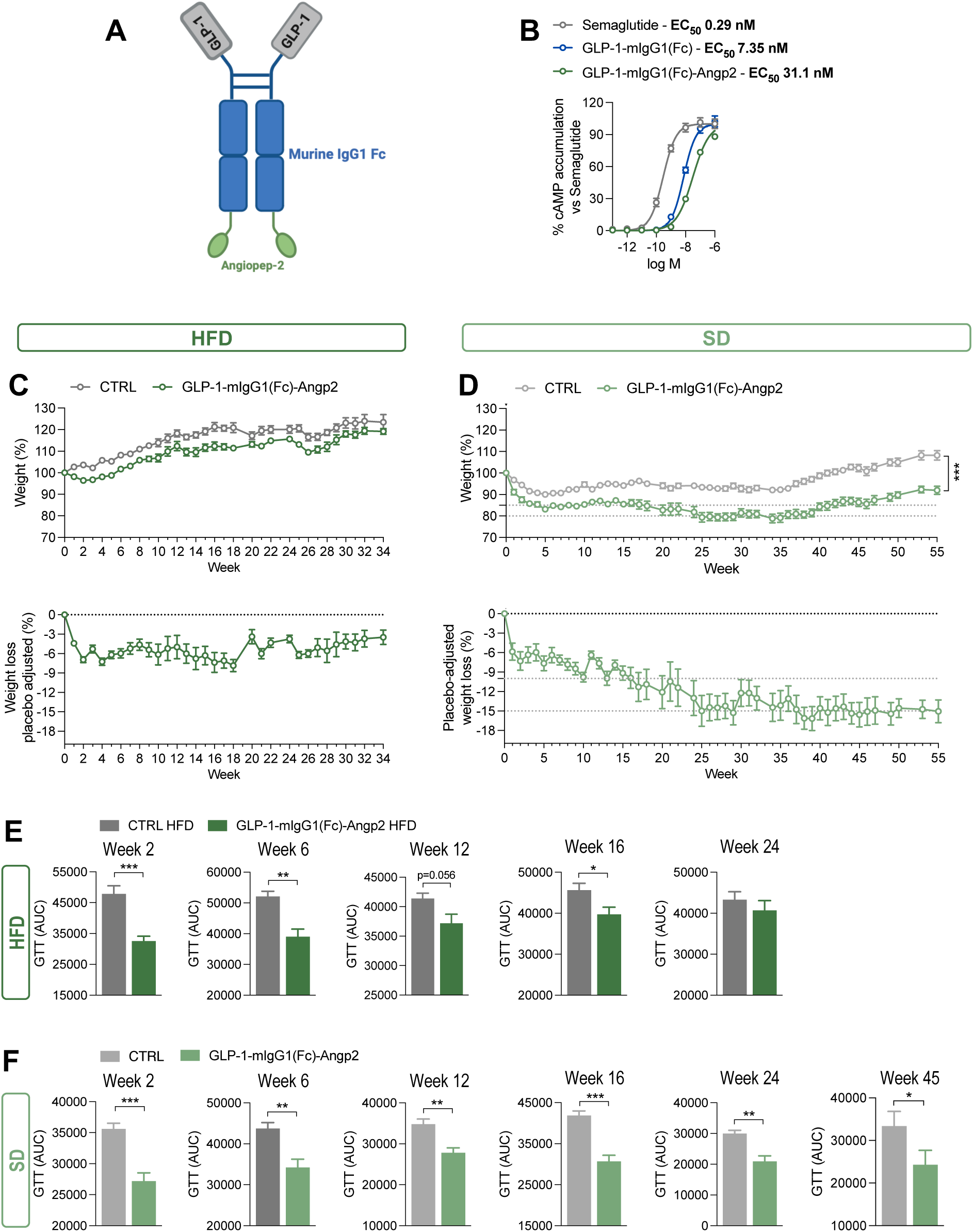
Enhanced BBB crossing dramatically improves GLP-1-mIgG1(Fc) efficacy. (A) Schematic of the protein. (B) Intracellular cAMP accumulation measured in HEK293T cells expressing GLP-1R. Data are expressed as % of cAMP relative to the highest semaglutide concentration (n≥4 from 3 independent experiments). Percentage of absolute (top panel) and placebo adjusted weight gain/loss (bottom panel) relative to the time of the EP (Week 0) in mice fed (C) high fat diet (HFD) (n=5), and (D) standard diet (SD) (n=5). Area under the curve (AUC) derived from glucose tolerance test (GTT) performed at the time points indicated in the figure in mice fed (E) HFD (n=5), or (F) SD (n=5). Data are expressed as mean ± SEM. *p<0.05, ** p<0.01, *** p<0.01. Statistical significance was determined by Two-way mixed-effects ANOVA (C-D) and unpaired t-test (E-F).

First, we evaluated the potency of GLP-1-mIgG1(Fc)-Angp2 on GLP-1R and compared it to that of semaglutide and GLP-1-mIgG1(Fc), using the in vitro reporter assay described earlier in this manuscript. While addition of Angiopep-2 to GLP-1-mIgG1(Fc) led to reduction of the potency, it remained at a low EC50 of 31.1 nM (Figure 4B). Thus, we decided to evaluate DNA-based delivery of this molecule in the DIO model, following the same diet protocol described in Figure 1A. Mice that remained on HFD lost ∼7% placebo adjusted weight in the first 2 weeks, and for the following 16 weeks they gained weight at a slower rate compared to controls (Figure 4C), but the weight started rebounding after week 18. Mice switched to SD, however, exhibited a much more robust therapeutic effect of GLP-1-mIgG1(Fc)-Angp2 after MYO delivery. They started losing weight immediately after IM-EP with ∼15% placebo adjusted weight reduction, sustained for over one year after a single administration (Figure 4D). We also assessed the response to glucose via GTT assay at different time points. Importantly, mice that received DNA-encoded GLP-1-mIgG1(Fc)-Angp2 showed an improved glucose clearance compared to the controls over an extended period (Figure 4E and F). The effect in animals on SD was more durable, and at week 45 still significantly differentiated from placebo. Interestingly, in the HFD cohort, by week 24 no difference in GTT was detected between treated and control animals, which correlated with weight rebound in the treated animals. These data suggest that targeting GLP-1-mIgG1(Fc) to the BBB, thereby enabling access to hypothalamic GLP-1R, is critical for sustained weight loss and long-term glucose control.

### Human Fc fusion demonstrates similar effects in an obesity model

Given the ultimate goal of developing a molecule suitable for clinical use, we replaced the murine IgG1 Fc domain with that of human IgG4, and retained the Angiopep-2 fusion. This molecule is referred to as GLP-1-hIgG4(Fc)-Angp2 (Figure 5A).

**Figure 5.**
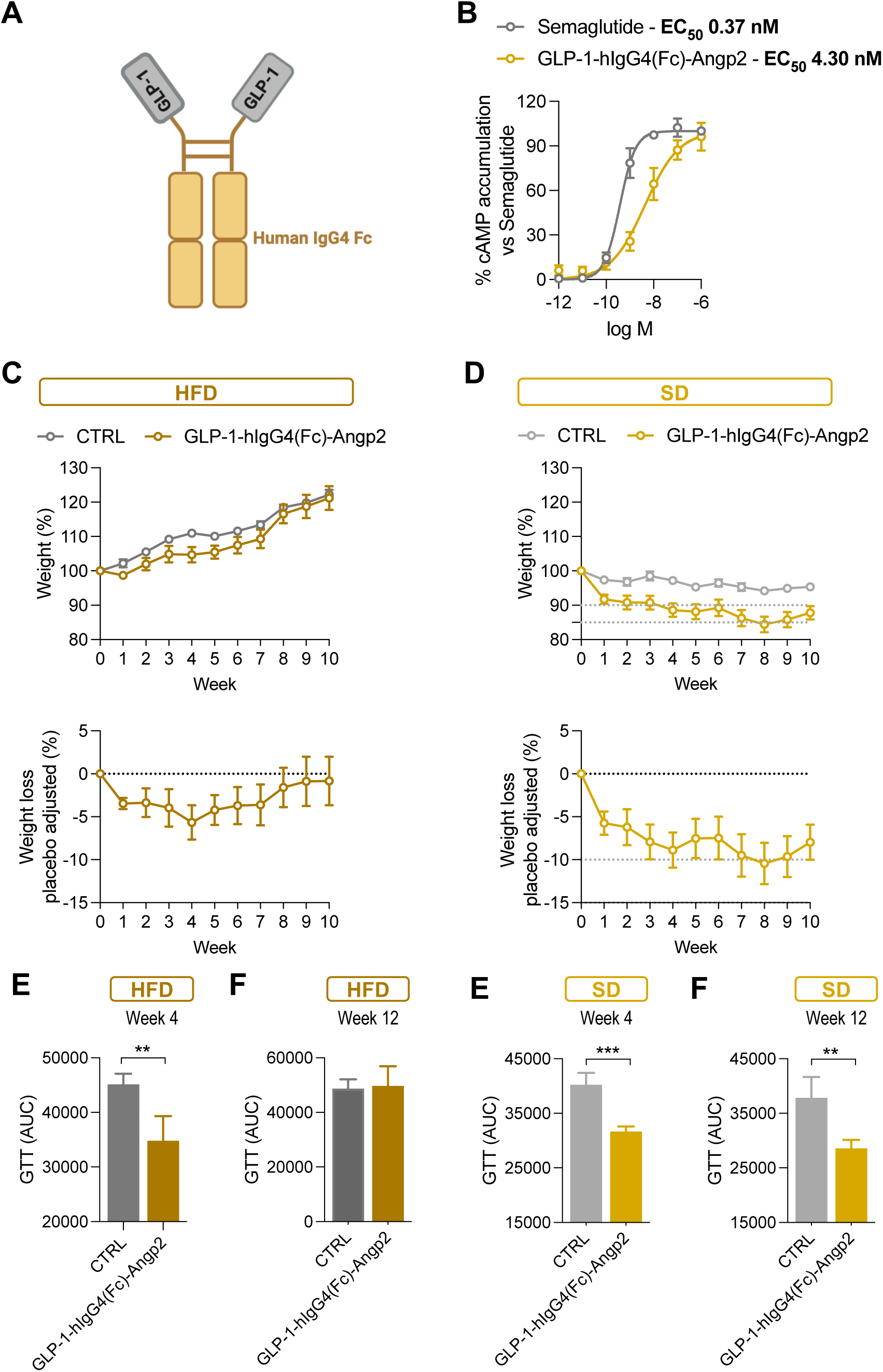
Human Fc fusion demonstrates similar effects in an obesity model. (A) Schematic of the protein. (B) Intracellular cAMP accumulation measured in HEK293T cells expressing GLP-1R. Data are expressed as % of cAMP relative to the highest semaglutide concentration (n=3). Percentage of absolute (top panel) and placebo adjusted weight gain/loss (bottom panel) relative to the time of the EP (Week 0) in mice fed (C) high fat diet (HFD) (n=5), and (D) standard diet (SD) (n=5). Area under the curve (AUC) derived from glucose tolerance test (GTT) performed at (E) week 4, and (F) week 12 in mice fed HFD (n=5), and at (G) week 4, and (H) week 12 in mice fed SD (n=5). Data are expressed as mean ± SEM. ** p<0.01, *** p<0.01. Statistical significance was determined by unpaired t-test (E-G).

First, we evaluated the potency of GLP-1-hIgG4(Fc)-Angp2 on GLP-1R and compared it to the potency of semaglutide, as described before. We determined that GLP-1-hIgG4(Fc)-Angp2 binds GLP-1R and exhibits low EC_50_ of 4.3 nM (Figure 5B).

We then evaluated the delivery of GLP-1-hIgG4(Fc)-Angp2 with MYO Technology in mice, following the same experimental design as before (Figure 1A). Mice that were administered DNA-encoded GLP-1-hIgG4(Fc)-Angp2 and kept on a HFD gained weight at a slower rate compared to controls, showing ∼7% placebo adjusted weight loss (Figure 5C). Mice that were switched to SD showed over 10% placebo adjusted weight loss (Figure 5D). Furthermore, GTT tests performed at week 4 showed a significantly improved glucose clearance compared to control, regardless of the diet (Figure 5E and 5G). The effect in animals on SD was maintained also at week 12, while in the HFD cohort, at week 12 no difference in GTT was detected between treated and control animals, which correlated with weight rebound in the treated animals (Figure 5F and 5H).

These data are consistent with our hypothesis that targeting a GLP-1Fc fusion to the brain via the addition of Angiopep-2 promotes a sustained weight loss and a better glucose clearance over time.

## MATERIAL AND METHODS

### Animals

All animal procedures were performed at Mispro facilities and overseen by the Mispro Institutional Animal Care and Use Committee (IACUC), with the animals cared for in accordance with all federal and state regulations. The mice were housed in disposable, individually ventilated, filter-top cages (Innocage®, Innovive Inc., San Diego, CA, USA). 6/8-week-old C57BL/6J (The Jackson Laboratory, strain #000664) male mice were fed high fat diet (HFD. ResearchDiet, cat# D12451) for 8 weeks. The day before study initiation all mice were weighed and randomized into treatment groups based on body weight (BW) so that all groups had similar starting BW before treatment (n = 5/group). Immediately after IM/EP, mice were either kept on HFD or switched to a standard diet (SD. ResearchDiet, cat# D12450H), and monitored as indicated in the result section. During the course of the study mice weight was recorded weekly. For the % of weight gain/loss (top panels), individual mouse weight recorded the day before IM/EP was considered 100%, and the weight at the following time-points calculated relative to this 100% baseline. For weight loss placebo adjusted, for each week the mouse average weight in the control cohort was considered 100%. Then, the individual weights in the treated cohort calculated as a percentage of this 100% baseline. The difference between this percentage and 100% represents the percentage of weight loss compared to the placebo group.

### Electroporation

Electroporation was performed bilaterally in the tibialis anterior (TA) muscle. Target muscle was injected with a 40 µL mixture consisting in 50 µg of pDNA and 3U of Hylenex® (hyaluronidase human injection, Halozyme Therapeutics), formulated in 0.45% saline, with pDNA at a final concentration of 1.25 μg/μL. This formulation was injected using a 29-gauge syringe (Becton Dickinson, cat# 324702), at a depth of 1.9 ± 0.2 mm. The pDNA injection was immediately followed by electroporation using an in-house-developed needle-based electroporator. The electroporator consists of four 0.3 mm diameter gold acupuncture needle electrodes (Asiamed, cat# AMG-3030), each at the corner of a 2 × 2 mm square. A series of pulses at 150 V/cm was applied using a RenBio custom-built generator (RenBio Inc).

### Glucose Tolerance Test (GTT)

After overnight fasting, basal glucose levels were recorded by using an Auvon glucose meter. Mice were then injected with a glucose (Sigma-Aldrich, cat# G7021) solution intraperitoneally (1.5 g/kg BW in water) and blood glucose measured at the indicated times.

### GLP-1R in vitro reporter assays

HEK293T (cat#) were seeded at 1.8e6 cells in a 10 cm plate in DMEM (Corning, cat# 10-017-CV) completed with 10% FBS (Corning, cat# 35-011-CV) and 1x pen/strep (Corning 30-002-CI). After 24h 2.5 ug of pDNA expressing GLP-1R and 2.5 ug of pGL4.29[*luc2P*/CRE/Hygro] (Promega, cat# E8471) were transiently transfected with PEI. DNA encoding GLP-1R (GenBank, Gene ID: 2740) was cloned into the pCBAINA, a gWiz-derived expression construct. The backbone of pCBAINA includes a cytomegalovirus (CMV) enhancer, chicken beta actin promoter, CMV intron A, bovine growth hormone polyadenylation (BGH-PolyA) sequence, and kanamycin resistance gene. GLP-1R sequence was synthesized by Twist Bioscience (San Francisco, CA, USA). Medium was changed 6h post-transfection to selection medium containing 0.5 ug/mL of puromycin (Boston Bioproducts, cat# ABT-440). 24h post-transfection cells were seeded at 70000/well in a 96 multiwell plate. 48h post-transfection cells were incubated with different concentrations of IRAs. After 3h of incubation cells were lysed with Bright-Glo™ Luciferase Assay System (Promega, cat# E2610) according to the manufacturer’s protocol, and the luminescence was measured using GloMax Navigator Microplate Luminometer (Promega, cat# GM2000). Luminescence data were normalized as % activation compared to the one of semaglutide at the maximum concentration.

## DISCUSSION

In recent years, the IRA class of drugs such as GLP-1 have transformed the treatment of obesity and T2D^3^. It is clear that these molecules not only have the potential to help patients lose weight, but also to make inroads in tackling many of the comorbidities associated with these diseases, including cardiovascular and kidney diseases^19–21^. A major drawback to these drugs, however, is their relatively short half-life, despite both genetic and biochemical modifications aimed at improving their pharmacokinetic (PK) profile. Because of this, the majority of the current generation of approved GLP-1 drugs must be dosed weekly by subcutaneous injection. And as a result of the expense associated with manufacturing protein biologics, as well as the frequent dosing regimen required, these drugs can exact a high cost from patients, often in the range of $1000 per month.

In order to address these barriers of frequent dosing and high costs, we have evaluated a DNA-based delivery approach. Previous work with our MYO Technology platform for the delivery of another small therapeutic protein with a short half-life, G-CSF, yielded very encouraging results regarding the ability of this approach to overcome the poor PK profile of some small proteins^6^ Given these results, we sought to determine if this approach could similarly improve the outcomes of treatment with IRAs.

We found that delivery of a GLP-1 RA alone could indeed lead to weight loss in a mouse DIO model. However, these effects were transient, and only significantly different from untreated controls when the animals were also switched to a standard diet following treatment. In order to probe the effect of modifications that would improve the half-life of these molecules, we generated and evaluated Fc-fusion variants, initially appending a murine IgG1 Fc. This Fc-fusion protein demonstrated a modest improvement in both weight loss and glucose metabolism as compared to the unmodified GLP-1 RA molecule.

Though questions remain about the various mechanisms by which the GLP-1 class of molecules exert their effects on weight loss, strong data indicate that targets in the brain play a role^1^. Thus, we sought to understand whether increased access to the brain, via improved penetration of the BBB, may lead to improved efficacy. Indeed, by appending Angiopep-2, a peptide known to increase BBB targeting^9^, we significantly improved the magnitude and duration of weight loss following a single administration, an effect that persisted for over one year. This was also accompanied by a durable improvement in glucose metabolism. Importantly, this effect was also apparent in the context of a human IgG1 fusion, though this study has not yet been followed as long.

These results clearly demonstrate that the addition of Angiopep-2 to a GLP-1 RA improves both the magnitude and durability of weight loss in a mouse DIO model, when delivered as a DNA-based drug with MYO Technology. In order to further probe the mechanism of this effect, we plan to evaluate the biodistribution of this new molecule, with a particular focus on the central nervous system.

Although the GLP-1 class of drugs has clearly demonstrated potency in terms of overall weight loss, improving the quality of weight loss remains one of the next important hurdles. Many groups are evaluating the ability of additional therapeutics to reduce lean muscle loss, primarily in the form of monoclonal antibodies targeting pathways involved in muscle maintenance and growth, such as myostatin or activin^22,23^. MYO Technology-based delivery is particularly well-suited to such multiplexing, in that delivery of multiple genes can be easily accomplished either on a single plasmid or via incorporation of multiple plasmids expressing different genes.

GLP-1 drugs have ignited a dramatic shift in obesity research and treatment, revealing both impressive efficacy and areas clearly needing improvement. We have demonstrated that a DNA-based delivery approach, using MYO Technology, has the potential to overcome the poor PK profile associated with these drugs. Additionally, by taking advantage of the ease of co-delivery of multiple biologics using this approach, we present an opportunity to evaluate many of the molecules aimed at improving the quality of weight loss achieved by these remarkable drugs.

## Notes

### Competing Interest Statement

The authors have declared no competing interest.

### Summary of Updates

Text modified to reflect the new results in figure 4

